# Orchard Management and Landscape Context Mediate the Floral Microbiome of Pear

**DOI:** 10.1101/2020.12.23.424173

**Authors:** Robert N. Schaeffer, Vera W. Pfeiffer, Saumik Basu, Matthew Brousil, Christopher Strohm, S. Tianna DuPont, Rachel L. Vannette, David W. Crowder

**Affiliations:** Department of Biology, Utah State University, Logan, UT 84322; Department of Entomology, Washington State University, Pullman, WA 99164; Tree Fruit Research and Extension Center, Washington State University, Wenatchee, WA 98801; Department of Entomology & Nematology, University of California Davis, Davis, CA 95616

**Keywords:** flower microbiome, integrated pest management, landscape heterogeneity, *Pyrus communis*

## Abstract

Crop-associated microbiota are key factors affecting host health and productivity. Most crops are grown within heterogeneous landscapes, and interactions between management practices and landscape context often affect plant and animal biodiversity in agroecosystems. However, whether these same factors typically affect crop-associated microbiota is less clear. Here, we assessed whether orchard management strategies and landscape context affected bacterial and fungal communities in pear (*Pyrus communis*) flowers. We found that bacteria and fungi responded differently to management schemes. Organically-certified orchards had higher fungal diversity in flowers than conventional or bio-based integrated pest management (IPM) orchards, but organic orchards had the lowest bacterial diversity. Orchard management scheme also best predicted the distribution of several important bacterial and fungal genera that either cause or suppress disease, with organic and bio-based IPM best explaining the distributions of bacterial and fungal genera, respectively. Moreover, patterns of bacterial and fungal diversity were affected by interactions between management, landscape context, and climate. When examining the similarity of bacterial and fungal communities across sites, both abundance- and taxa-related turnover were mediated primarily by orchard management scheme and landscape context, and specifically the amount of land in cultivation. Our study reveals local- and landscape-level drivers of floral microbiome structure in a major fruit crop, providing insights that can inform microbiome management to promote host health and high-yielding quality fruit.

**IMPORTANCE:** In tree fruits, proper crop management during bloom is essential for producing disease-free fruit. Tree fruits are often grown in heterogeneous landscapes; however, few studies have assessed whether landscape context and crop management affect the floral microbiome, which plays a critical role in shaping plant health and disease tolerance. Such work is key for identification of tactics and/or contexts where beneficial microbes proliferate, and pathogenic microbes are limited. Here, we characterize the floral microbiome of pear crops in Washington State, USA, where major production occurs in inter-mountain valleys and basins with variable elevation and microclimates. Our results show that both local (crop management) and landscape (habitat types and climate) level factors affect floral microbiota, but in disparate ways for each kingdom, suggesting a need for unique management strategies for each group. More broadly, these findings can potentially inform microbiome management in orchards for promotion of host health and high-quality yields.

## INTRODUCTION

Microbial communities affect plant health and productivity. For agricultural crops, microbes can affect nutrient mobilization and transport, often promoting plant growth and disease resistance (Pii *et al*., 2015; Vurukonda *et al*., 2016; Berg and Koskella, 2018). In turn, understanding and managing microbiome assembly could enhance agricultural sustainability by reducing reliance on external inputs, enhancing yields, and potentially contributing to the maintenance of both biodiversity and the functioning of agricultural landscapes (Mueller and Sachs, 2015; Busby *et al*., 2017; Toju *et al*., 2018). Yet, despite the growing recognition of the importance of the microbiome to crop productivity, processes governing the assembly of microbiomes for many crop species are still largely unclear (but see Edwards *et al*., 2015; Grady *et al*., 2019).

Agricultural landscapes are often spatially heterogeneous. Accruing through shifts in land tenure over time, this heterogeneity reflects a landscape’s composition and configuration (Fahrig and Nuttle, 2005; Fahrig *et al*., 2011; Smith *et al*., 2020). Specifically, crop production occurs on patches of land that exist within habitat mosaics containing patches of the same crop, alternative commodities, and semi-natural vegetation. Such variation in land cover around a crop field may strongly affect local abiotic and biotic conditions. Most studies assessing the effects of spatial context, however, have focused primarily on plants (Smith *et al*., 2020) and animals (Karp *et al*., 2018), but effects of landscape-level drivers on plant-associated microbiomes has received less attention. This is a problematic knowledge gap as microbes often disperse over long distances, and studies show that spillover of microbes from agricultural into natural habitats is affected by landscape context and dispersal ability of individual taxa (Bell and Tylianakis, 2016). Many microbes are often affected strongly by environmental conditions, and abiotic variation across landscapes can sometimes predict outbreaks of pathogenic microbes (Smith and Pusey 2010)

At the orchard scale, management practices employed to control pests and disease can also shape microbiome assembly and structure. Agricultural producers often rely on agrochemicals to prevent establishment or directly suppress both pests and pathogens. As part of an integrated pest management (IPM) program, these practices can vary in intensity across orchards, including the frequency of application, the active ingredients of chemical controls, and how they are coupled with other biological or cultural-control strategies (Agrios, 2005). Indeed, the application of antibiotics, fungicides, or microbiological control agents can leave distinct signatures on the microbiome associated with tree fruits (Johnson and Stockwell, 1998; Schaeffer *et al*., 2017). Though their application can often have direct, suppressive effects on the abundance of targeted, pathogenic taxa (Johnson and Stockwell, 1998), non-target effects on associated yeasts and bacteria have also been observed (McGhee and Sundin, 2011; Schaeffer *et al*., 2017).

Here, we assessed how local- and landscape-level processes affected the diversity and structure of microbe communities associated with pear (*Pyrus communis*) flowers in Washington State, USA. We focused on microbes on flowers, as these ephemeral structures produce the fruit, but are also the primary infection site for pathogens such as the bacterium *Erwinia amylovora*, the causal agent of fire blight (Vanneste 2000). As a consequence, pear orchards are typically heavily managed during bloom to minimize disease risk while promoting pollination (McGregor 1976, Johnson and Stockwell 1998). Such management tactics range from the use of managed honey bees, to application of diverse bactericides for control of fire blight. We predicted that floral microbiota would be impacted by orchard management practices and the abiotic and biotic landscape conditions. Such work provides important insights into microbial colonization and community structure pre- and post-pollination, important windows for production.

## RESULTS

### Pear flower microbiome

Our study sampled bacterial and fungal communities associated with pear flowers across 15 orchards with three management types (conventional, bio-based IPM, and organic; 5 sites of each). After quality-filtering and processing, we detected 142 bacterial and 1703 fungal amplicon sequence variants (ASVs) from the pear flowers. The bacterial community was dominated by members of the *Bacillaceae, Enterobacteriaceae, Lactobacillaceae*, and *Pseudomonadaceae* (Fig. 1A), with each family comprising on average, 22%, 15%, 9%, and 9% of sequences, respectively. Beneficial bacteria previously found to be associated with disease suppression in this system (i.e., *Bacillus, Pantoea*, and *Pseudomonas*) comprised ~11% of taxa (ASVs) observed, and ~41% of the relative abundance. The fungal community was dominated by members of *Aureobasidiaceae*, *Cladosporiaceae*, *Mycosphaerellaceae*, and *Sclerotiniaceae* (Fig. 1B), with each family comprising on average, 16%, 8%, 14%, and 7% of sequences, respectively. Of the *Aureobasidiaceae*, four ASVs were identified to the species level as *Aureobasidium pullulans*, a beneficial fungus used for biological control of fire blight. Twenty-one additional ASVs were identified as belonging to genera *Botrytis, Cladosporium, Monilinia, Mycosphaerella*, or *Penicillium*, potentially important agents of pre- and post-harvest disease.

**Figure 1.**
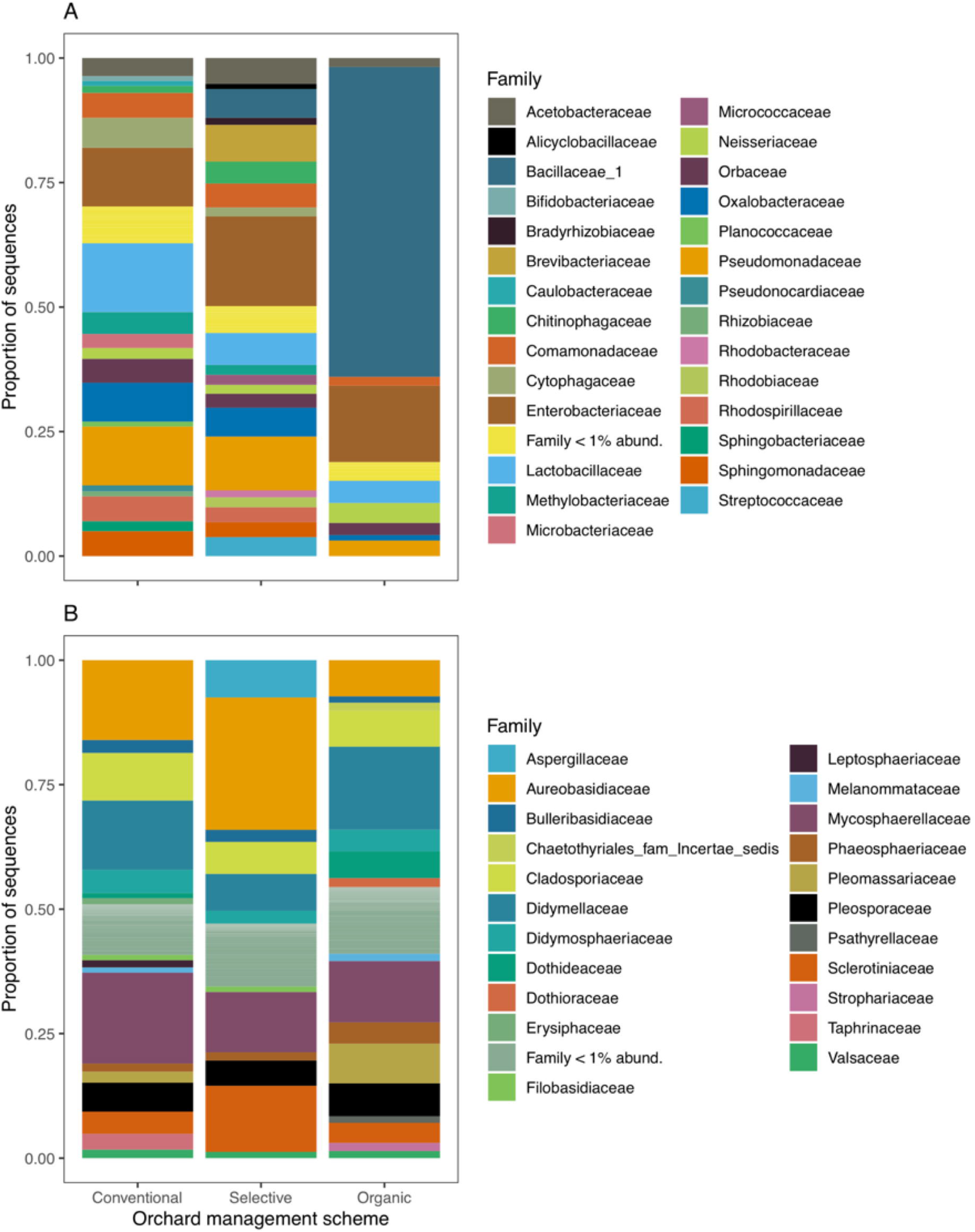
Relative abundance (Proportion of sequences) of (A) bacterial and (B) fungal families associated with pear flowers. Flowers were collected from orchards that reflected three unique management schemes (Conventional, bIPM, Organic).

### Orchard management and landscape context affect bacterial and fungal alpha diversity

Orchard pest management practices were significantly associated with pear flower bacterial and fungal diversity (Shannon Index) (Table 1). Considered alone, conventional and bIPM-managed orchards were found to have ~60% higher bacterial diversity than those managed organically (Fig. 2A), while organically managed orchards exhibited the highest fungal diversity (Fig. 2B). Yet, the positive effects of organic management on fungal diversity were not significant in the multiple variate linear model when controlling for land cover and climate. In these linear regression models, both organic and bIPM-management styles reduced bacterial and fungal diversity, although the negative influence of organic management on fungal diversity was weak. Land cover was also associated with bacterial and fungal diversity: bacterial diversity declined with increasing proportion of habitat containing forest or pear, while fungal diversity increased with pear crop cover. Microclimatic conditions were also associated with both bacterial and fungal diversity, though minimum temperature was the only variable of significant effect on fungi, and minimum VPD was for bacteria in the top AIC selected model (Table 1).

**Figure 2.**
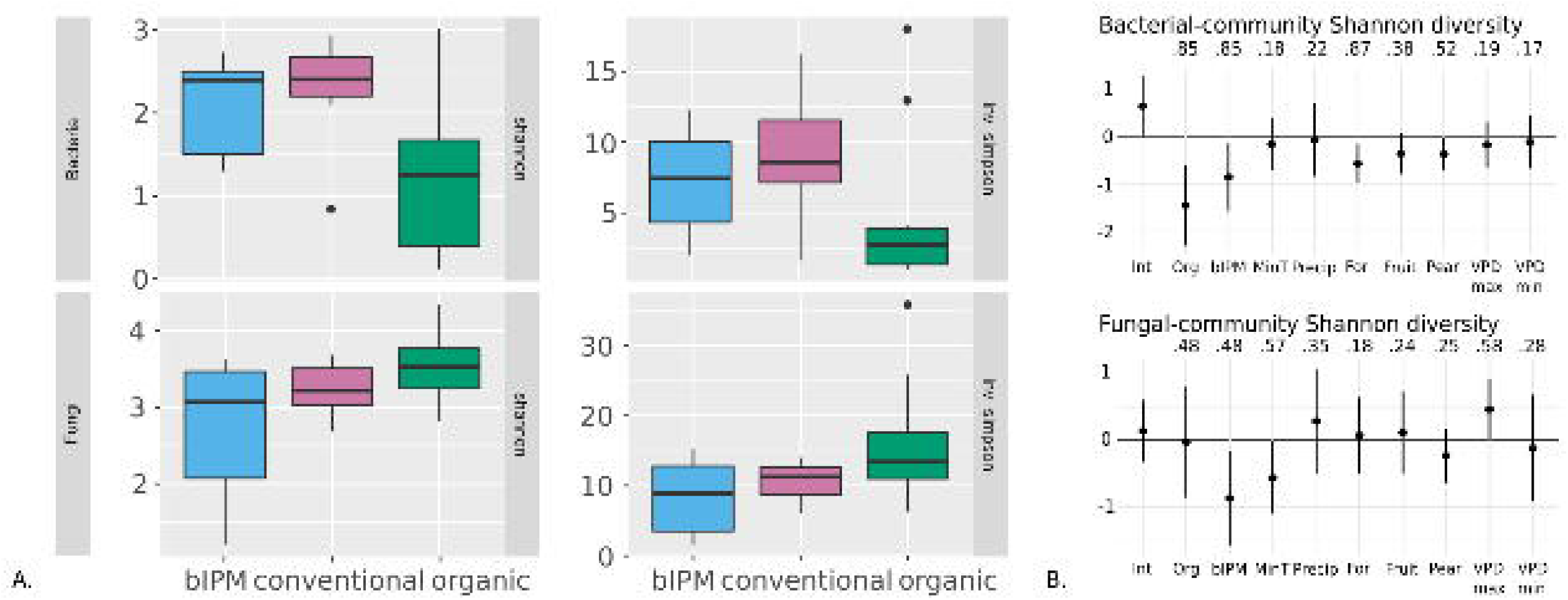
(A) Boxplots of Shannon diversity by orchard management style and (B) coefficients from the 90% confidence set of top multivariate models. Variable importance was evaluated as the number of models within the 90% confidence model set in which the factor was included.

**Table 1.** Multivariate linear regression models for bacterial and fungal Shannon diversity. Top models were selected by AICc.

| Bacteria |  |  |  |  |  |
| --- | --- | --- | --- | --- | --- |
| Variable | Estimate | Std Error | *P* | Model *adj R*<sup>2</sup> | *P* |
| Intercept | 0.869 | 0.274 | 0.005 | 0.522 | <0.001 |
| Organic management | -1.728 | 0.478 | 0.002 |  |  |
| bIPM<sup>a</sup> management | -0.964 | 0.369 | 0.016 |  |  |
| Proportion of landscape - forest | -0.597 | 0.186 | 0.004 |  |  |
| Proportion of landscape - pear | -0.506 | 0.187 | 0.013 |  |  |
| VPD<sup>b</sup> minimum | -0.498 | 0.281 | 0.090 |  |  |
| VPD maximum | -0.411 | 0.250 | 0.114 |  |  |
| Fungi |  |  |  |  |  |
| Variable | Estimate | Std Error | *P* | Model *adj R*<sup>2</sup> | *P* |
| Intercept | 0.460 | 0.294 | 0.131 | 0.426 | 0.002 |
| Organic management | -0.357 | 0.494 | 0.477 |  |  |
| bIPM management | -1.022 | 0.392 | 0.015 |  |  |
| Proportion of landscape - pear | 0.367 | 0.153 | 0.025 |  |  |
| VPD maximum | -0.252 | 0.188 | 0.191 |  |  |
| Minimum temperature | -0.411 | 0.184 | 0.035 |  |  |

<sup>a</sup>Biological-based Integrated Pest Management
<sup>b</sup>Vapor Pressure Deficit

### Orchard management practices drive the distribution of pathogenic fungal species and the presence of bacterial genera associated with disease suppression

Focal bacterial and fungal genera of concern were first investigated to assess the scale of spatial autocorrelation, as well as potential associations with aspects of landscape context. Positive spatial autocorrelation was exhibited for each of the nine taxa examined, but only at the shortest distances of less than 1 km. Using canonical correlation analysis to assess how landscape and management variables were associated with the microbial species composition, we found significant associations between predictors and bacterial (Table S1; Pillai’s trace *P* = 0.014) and fungal communities (Table S2; Pillai’s trace *P* = 0.005) (Fig. 3). The three bacterial genera associated with disease suppression were distributed very differently in association with the factors of interest. More specifically, the relative abundance of *Bacillus*, a bacteria commonly applied to suppress disease in pear, was most strongly associated with organic management (Fig. S1), followed closely by the amount of surrounding forest, and then geographic distance. These top factors, aligned with Axis 1, were negatively associated with *Pseudomonas*, while bIPM was the most important predictor of *Pantoea* (more aligned with Axis 2). Similar to *Pantoea*, bIPM (+) and organic management (−) best predicted the presence of *Aureobasidium*, a beneficial fungi aligned with Axis 1, and *Monilinia* to a lesser degree. Minimum temperature (+) and minimum VPD (+) best predicted *Botrytis*, *Cladosporium*, and *Mycosphaerella*, as well as *Monilinia* (−), all pathogenic fungi of concern for pears. Finally, the proportion of forest in the landscape, and geographic distance, were associated with the distribution of these fungal genera of interest (Table S2).

**Figure 3.**
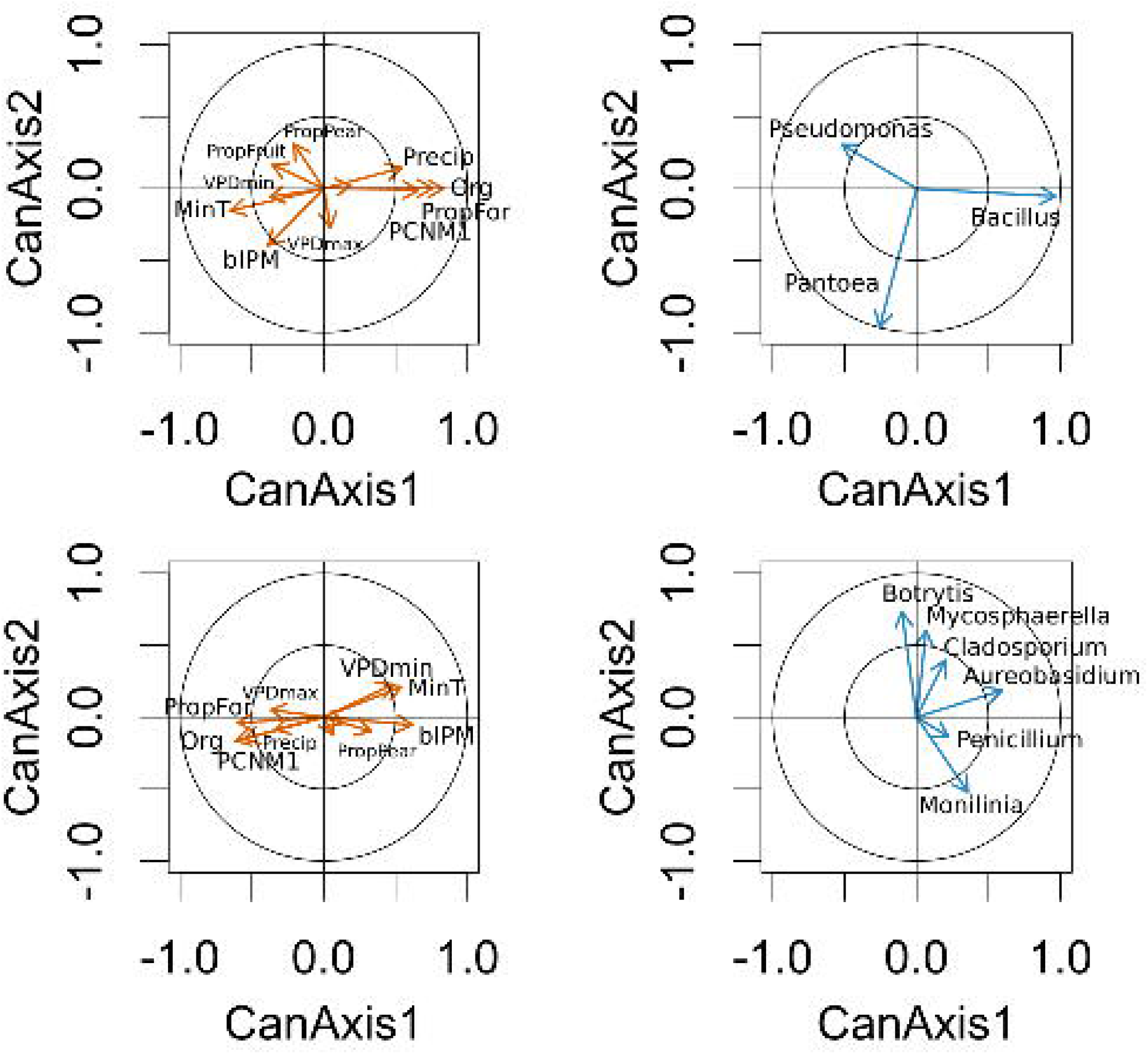
Canonical correlation analysis of three beneficial bacteria taxa and five pathogenic fungal taxa. The left panel depicts the variance explained by the factors in the canonical axes, and the right panel depicts the variance explained by the canonical axes in the taxa of interest.

### Microbial beta diversity was affected by orchard management and landscape context

Overall bacterial community similarity, and turnover of specific taxa across orchards, was best predicted by geographic distance between orchards and orchard management (Fig. 4; Table 2). In other words, sites that were located nearby, or had the same management scheme, tended to be most similar in terms of community composition. In contrast, abundance-related turnover across sites was affected mainly by the proportion of landscape under fruit cultivation, namely apple. With respect to fungi (Fig. 4; Table 3), turnover of fungal communities across sites was associated with the amount of pear production in the landscape, temperature, and vapor pressure deficit (VPD). Temperature and VPD, along with surrounding forest, were important drivers of taxa-related turnover. In contrast, abundance-related community turnover was associated with geographic distance and the proportion of landscape represented by forest around orchards.

**Figure 4.**
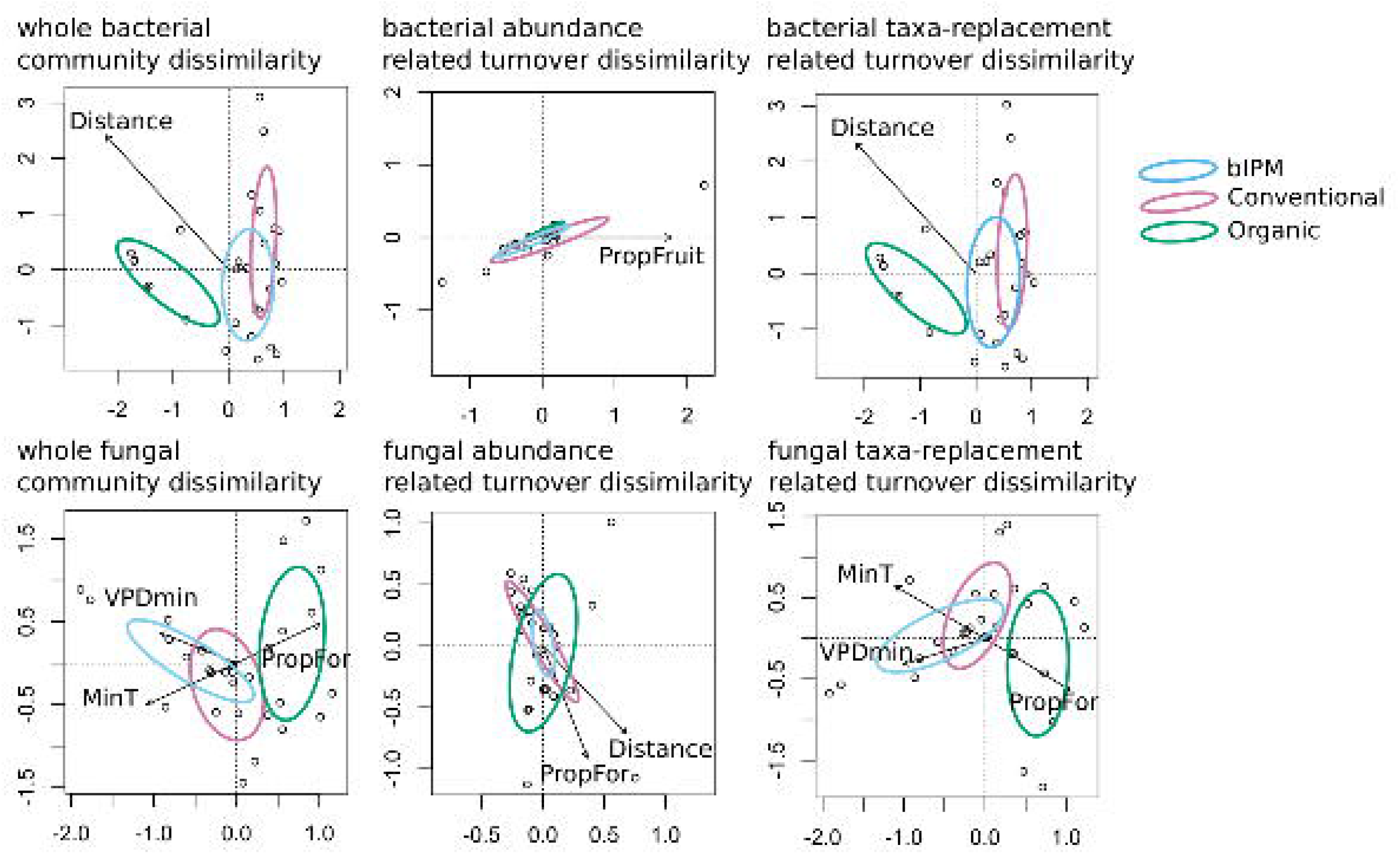
Restricted Distance-based Analysis of bacterial and fungal community beta diversity and explanatory variables included in the top AIC-selected RDA models. Variance explained by each factor is in Tables 2 and 3.

**Table 2.** Results from Restricted Distance-based Analysis (RDA) of bacterial community beta-diversity. Top model selected by AIC.

| Whole community beta-diversity |  |  |
| --- | --- | --- |
| Variable | *adj R*<sup>2</sup> | Pr (>*F*) |
| Orchard management scheme | 0.147 | 0.002 |
| Geographic distance | 0.172 | 0.026 |
| Abundance-related community beta-diversity |  |  |
| Proportion of landscape - fruit | 0.594 | 0.040 |
| Taxa-related community beta-diversity |  |  |
| Orchard management scheme | 0.164 | 0.002 |
| Geographic distance | 0.195 | 0.026 |

**Table 3.** Results from Restricted Distance-based Analysis (RDA) of fungal community beta-diversity. Top model selected by AIC.

| Whole community beta-diversity |  |  |
| --- | --- | --- |
| Variable | adj $R^2$ | Pr ($>F$) |
| Proportion of landscape - pear | 0.170 | 0.046 |
| Minimum temperature | 0.092 | 0.002 |
| VPD<sup>a</sup> minimum | 0.132 | 0.018 |
| Abundance-related community beta-diversity |  |  |
| Geographic distance | 0.266 | 0.008 |
| Proportion of landscape - forest | 0.579 | 0.002 |
| Taxa-related community beta-diversity |  |  |
| Proportion of landscape - forest | 0.177 | 0.046 |
| Minimum temperature | 0.089 | 0.002 |
| VPD minimum | 0.134 | 0.026 |

<sup>a</sup>Vapor Pressure Deficit

## DISCUSSION

The Pacific Northwest is responsible for ~80% of pear production in the United States (USDA NASS 2019). Pre- and post-harvest diseases that can take hold during bloom threaten production and the quality of yield, however. Here, we investigated how local, orchard-level IPM practices interacted with landscape-level growing conditions to influence the structure and diversity of microbiota associated with pear flowers, potential sites for infection. Our analyses revealed that orchard management scheme can significantly influence the structure and diversity of both bacterial and fungal communities. Beyond local, orchard-level management, land cover and climate were also found to be significant predictors of microbe diversity, and bacterial and fungal communities were sensitive to different habitat types found in landscapes surrounding orchards. Finally, fungal alpha and beta diversity were far more sensitive than bacteria to microclimatic conditions experienced in orchards. In the sections that follow, we discuss these findings in light of understanding the key drivers of floral microbiome structure in this system.

### Orchard management mediates microbial diversity

Bacterial and fungal alpha diversity responded differently to orchard management scheme. Bacterial diversity was significantly higher in conventional and bIPM orchards compared to organic orchards; however, the opposite pattern was observed for fungi. Organic orchards had a high relative abundance of *Bacillus*, likely because of its application as a biological control agent. The strong effect of orchard management on bacterial diversity suggests that application of *Bacillus* reduced bacterial diversity, which may occur through resource competition, priority effects, or mass effects. *Bacillus* species have shown promise in limiting the establishment and development of the bacterial pathogen *E. amylovora*, the causal agent of fire blight (Sundin *et al*., 2009; Shemshura *et al*., 2020), and may also affect other floral microbes. Indeed, increased fungal diversity in organically-managed orchards could be a consequence of *Bacillus* application, although we were unable to directly assess if fungal abundance was affected in our study. In contrast to bacteria applied for biological control, we observed that *Aureobasidium* had a higher relative abundance in conventional and bIPM orchards than organic ones (where it was applied in one orchard for biological control). Background levels of some microbial taxa may be high and more prevalent in the presence of particular landscape and climate conditions (e.g., higher precipitation and high proportion of forest; Tables S1 and S2). These patterns may represent preferential use of these biological treatments across orchards in our survey. Though unable to confirm whether ASVs recovered in our dataset are these exact commercial strains, biologicals applied to pear flowers often have a high recovery rate in surveys (Stockwell *et al*., 2002; Johnson and Temple, 2013).

### Land cover and microclimate shape microbial diversity

Our results show that habitat patches with alternate tree fruit crops (apple, cherry) were negatively associated with both bacterial and fungal diversity on pear flowers, and appeared to be primary drivers of microbial community structure (Tables 2–3). Pear orchards in the Wenatchee River Valley are primary located in narrow, inter-mountain areas with highly variable elevation and land cover, including forest, additional pear orchards, and those dedicated to production of other deciduous fruits, namely apple. Vegetation in and around orchards can be an important source of inocula via airborne dispersal (Lindow and Andersen, 1996; Lymperopoulou *et al*., 2016). Furthermore, previous work on apple and pear flowers has revealed considerable overlap in the identity of microbes associated with each host species (Stockwell *et al*., 1999; Pusey *et al*., 2009; Smessaert *et al*., 2019). Such overlap, in addition to a reduction in diversity with increasing land cultivation, suggests a role for several key processes in shaping floral microbiomes in tree fruits. First, there is a high degree of shared usage of disease and pest management practices employed in pear and apple production systems, as both can suffer greatly from fire blight disease. Inputs applied in conventional and bIPM orchards, including antibiotics and fungicides (Table S3) can act as strong environmental filters on potential floral colonists (McGhee and Sundin, 2011; Schaeffer *et al*, 2017), or serve as a source for inocula when applied as biologicals, as observed in organic orchards. Second, both apple and pear systems rely considerably on honey bees (*Apis mellifera*) for pollination, which are known to leave a distinct imprint on floral microbiome diversity (Aizenberg-Gershtein *et al*., 2013). Increased reliance on a single pollinator species, combined with chemical and non-chemical inputs, are likely important contributors to patterns observed.

### Bacterial and fungal community turnover and dispersal

Orchard management scheme was a key determinant of bacterial community similarity across sites; however, other predictors often explained high levels of variance in community structure across sites. In particular, geographic distance explained a significant amount of variance in both whole-community and taxa-related beta-diversity of bacteria. In contrast, for fungi, spatial distance was a significant predictor of only abundance-related turnover. Beyond distance, climatic conditions contributed significantly to explained variance in the beta-diversity of fungal communities. In particular, VPD and temperature were negatively associated with fungal diversity, suggesting both microclimate variables affecting either species-specific patterns of growth and/or competition. Moisture availability is also an important determinant of microbial growth on the surface of plant tissues (Beattie 2002), with free water and humidity often being necessary for conidial germination, germ tube growth, and potential penetration of plant tissues, including floral organs. This has been frequently observed in other flowering systems of commercial value, including blueberries (Ngugi and Scherm, 2004), raspberries (McNicol *et al*., 1985), strawberries (Bulger *et al*., 1987), and cut roses (Muñoz *et al*., 2019). Within these systems, infection of the gynoecium can be a primary route of disease development. Alternatively, infection of petals and other organs can facilitate secondary infections of fruits (Petrasch *et al*., 2019). Of the fungal genera examined in our study, *Botrytis* has been documented to successfully infect the mesocarp via stamen filaments (de Kock and Holz, 1992). For the others of interest, it is unclear if there is a link between flower colonization and resulting development and pre- and post-harvest diseases.

More broadly, our results provide insight into local- and landscape-level drivers of floral microbiome diversity in an important tree fruit commodity, pear. Given the critical link between flowers, yield, and disease, identifying such drivers across both spatial and temporal scales could improve the understanding of links between management, host microbiome structure, and potentially disease resistance or susceptibility. With growing appreciation for the role of host microbiota in affecting resistance against disease (Berg and Koskella, 2018; Vannier *et al*., 2019), such information has potential to inform development of sustainable management practices in many different types of agroecosystems.

## MATERIALS AND METHODS

### Landscape survey

We surveyed 15 orchards throughout the Wenatchee River Valley of central Washington, USA (Fig. 5) in spring 2018. Within the United States, Washington State is the leading producer of deciduous tree fruit crops such as apples, pears, and cherries. These, as well as other commodities, are grown in variable inter-mountain river valleys and basins east of the Cascade mountains. These production areas generally experience temperate, dry conditions, in addition to favorable access to irrigation water originating from streams and rivers fed by snowmelt (Smith, 2000). Given the diverse topography of this region, however, individual orchards range in elevation from 20 to 1000 m above sea level (Smith, 2000). Key stages of fruit production, such as flower bloom, can thus experience considerable variation in microclimatic conditions among orchards, affecting bloom timing, fertilization, and fruit development (Logan *et al*., 2000; Lopez and DeJong, 2007). As flowers are habitat for diverse microbiota (Vannette, 2020), including a number of pathogenic species that cause pre- and post-harvest diseases of tree fruits (Ngugi and Scherm, 2006), microclimatic conditions could affect habitat quality, as well as colonization dynamics and the resulting structure of the floral microbiome.

**Figure 5.**
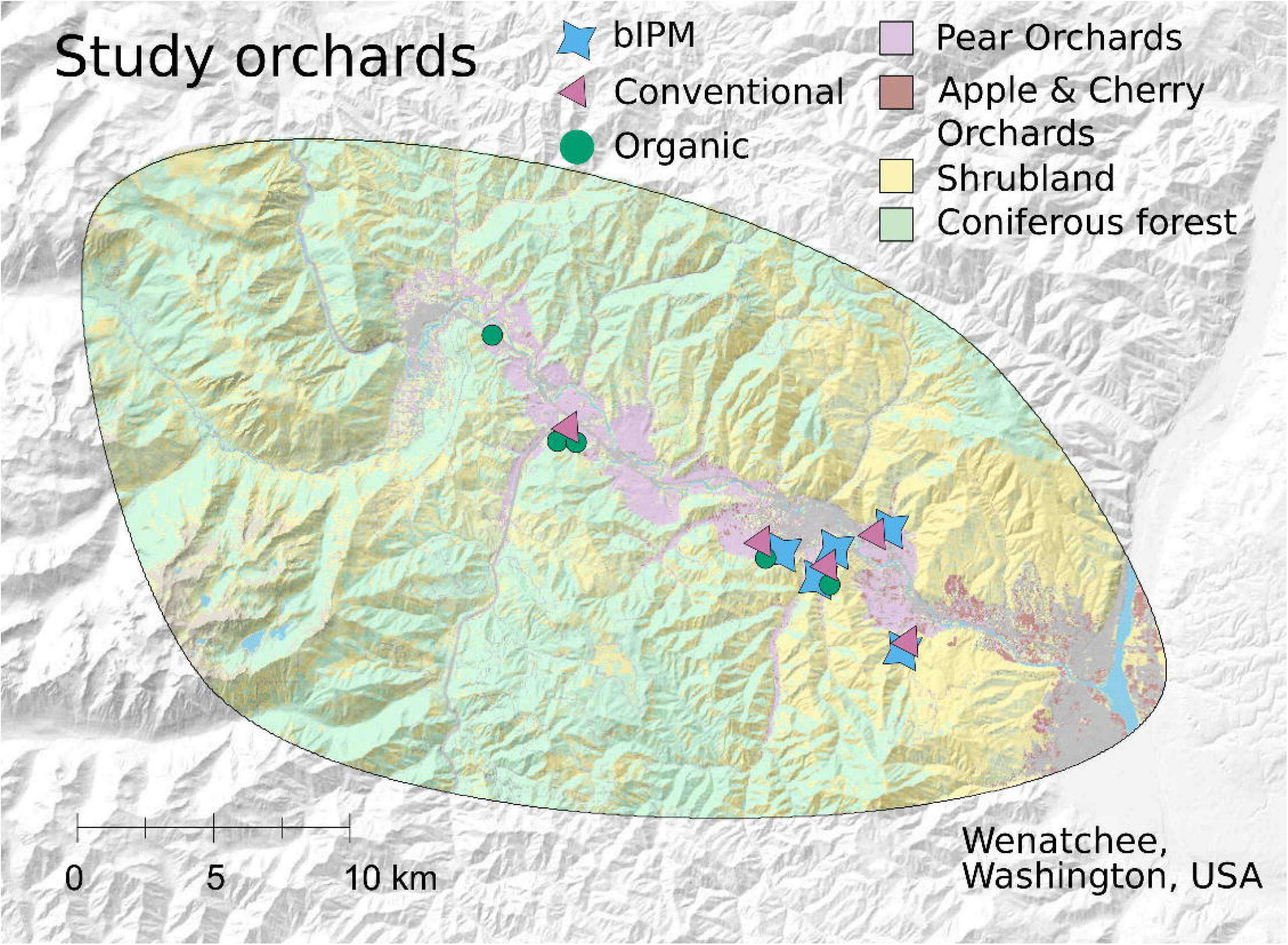
Geographic extent of survey, where fifteen pear orchards in central Washington across variable landscape contexts were sampled during peak bloom.

Our survey assess microbe communities on orchards that used one of three management schemes, with five replicates per scheme: organically-certified, conventional, and biological based integrated pest management (bIPM) (DuPont and Strohm, 2020). With each of these broad management types, growers were not restricted to a specific spray schedule, but each used a defined set of tools for pest and disease management (Table S3; DuPont and Strohm, 2020). Conventional management followed a standard practice (e.g., application of synthetic pesticides), while organic orchards were all managed following USDA-certified organic standards, which prohibits use of such synthetic chemicals. To control fire blight, organic producers often use Serenade^®^ Opti (Bayer CropScience, St. Louis, MO, USA) at full bloom, a bio-based fungicide and bactericide that leverages *Bacillus subtilis* (strain QST 713) endospores and its metabolic by products as active ingredients (DuPont *et al*., 2019). Serenade^®^ is not the only bio-based product leveraged by producers for control of fire blight in pear, however, and other products such as Blossom Protect™ (Westbridge Agricultural Products, Vista, CA, USA) can be used across organic, bIPM, and conventional schemes. Blossom Protect™ is derived from air-dried spores of *Aureobasidium pullulans* (strains DSM 14940 and 14941), an epiphytic or endophytic fungus associated with a wide range of plant species, including many tree fruits (Kunz 2006, Kunz *et al*. 2008, Leibinger *et al*. 1997). For those orchards that employed the bIPM scheme, growers used a toolbox of cultural controls combined with pesticides with less documented negative impact on natural enemies and other beneficial organisms. Such products included lime sulfur, kaolin, spinosad, and biologicals, applied at various stages of bloom stage (DuPont and Strohm, 2020).

Orchards were sampled once at peak bloom, either on April 30^th^ or May 1^st^ of 2018. At each orchard, 10 trees (‘Bradford’ variety) were sampled: five near the edge of the orchard and five in the interior. We chose this approach because previous studies suggest that semi-natural habitat in the surrounding landscape can both support and increase rates of visitation by native pollinators such as bees and flies (Klein *et al*., 2012). Moreover, pollinators can be important dispersal agents for microbes (Aizenberg-Gershtein *et al*., 2013; Vannette and Fukami, 2017); thus, our aim was to detect potential contributions of pollinator visitation to flower microbiome assembly in orchards. For each site (i.e., edge or interior) and sampling event, 50 open flowers (*N* = 10 per tree) were collected using aseptic technique and pooled at the site level. Flowers with flat, fully-reflexed petals that had been open for ~3 days were collected. Once collected, flowers were placed in a cooler and transferred to the lab, then stored at 4°C until processing.

### Sample processing

In a laboratory, whole flowers were washed with 20 mL of 1x-0.15% PBS-Tween solution, and samples were sonicated for 10 min to dislodge epiphytic microbes. After sonication, floral tissue debris was removed from sample tubes by pouring samples through autoclaved cheesecloth into a new, sterile Falcon tube. Falcon tubes containing debris-filtered samples were centrifuged at 3000 rpm for 10 min at 4°C to pellet microbial cells. We poured off the supernatant, re-suspended microbial cell pellets in 1 mL of autoclaved PBS solution, vortexed tubes, then transferred the cell suspensions to new 1.7 mL microcentrifuge tubes.

### DNA extraction and sequencing

Genomic DNA was extracted from samples using a ZymoBIOMICS^®^ DNA Microprep kit (Zymo Research, Irvine, CA, USA) following the manufacturer’s protocol. Extracted DNA was then used as template for library preparation and amplicon sequencing following Comeau et al. (2017), performed at the Centre for Comparative Genomics and Evolutionary Bioinformatics at Dalhousie University (Halifax, Nova Scotia, Canada). There, amplicon fragments were PCR-amplified from DNA in duplicate, using separate template dilutions (1:1 & 1:10) and high-fidelity Phusion polymerase (New England BioLabs Inc., Ipswich, MA, USA). A single round of PCR was performed using “fusion primers” (Illumina adaptors + indices + specific regions) targeting either the 16S V4-V5 (Bacteria; Primers: 515FB and 926R; Parada *et al*. 2015; Walters *et al*. 2015) or ITS2 (Fungi; Primers: ITS86 and ITS4; Op De Beeck *et al*. 2014) regions with multiplexing. PCR products were verified visually by running a high-throughput Invitrogen 96-well E-gel (Thermo Fisher Scientific Corp., Carlsbad, CA, USA). Any samples with failed PCRs (or spurious bands) were re-amplified by optimizing PCR conditions to produce correct bands to complete a sample plate before continuing with sequencing. The PCR reactions from the same samples were pooled in one plate, cleaned, and then normalized using the high-throughput Invitrogen SequalPrep 96-well Plate Kit (Thermo Fisher Scientific Corp.). Samples were then pooled to make one library, then quantified fluorometrically before sequencing. Amplicon samples were then run on an Illumina MiSeq using 300+300 bp paired-end V3 chemistry. Raw sequences are available on the NCBI Short Read Archive (SRA) under BioProject PRJNA659266.

Demultiplexed sequences were trimmed of trailing low-quality bases using the *DADA2* pipeline (v.1.8.0; Callahan *et al*., 2016) in R (v. 3.5.2; R Core Team, 2013). Paired-end reads were then quality-filtered, error-corrected, and assembled into ASVs. Once assembled, chimeras were detected, removed, and taxonomic information was then assigned to each ASV using the RDP Naïve Bayesian Classifier (Wang *et al*., 2007) trained to either the RDP training set (v.14) or UNITE general fasta release (v.7.2) for bacteria or fungi respectively. ASVs that failed to classify to kingdom, or identified as chloroplast or mitochondrial sequences, were discarded. Further, potential contaminant ASVs were identified through inclusion of negative controls during sample and sequence processing, and then removed using the ‘prevalence’ method with the *decontam* package in R (Davis *et al*., 2018). This filtering resulted in samples sequenced at a mean depth of 43,057 sequences per sample for bacteria and 25,890 for fungi. Samples were then rarefied (bacteria: 49; fungi: 14,920), with all but one bacterial sample (19orgedge) retained in the analyses that follow. Such a low cutoff for bacteria is consequence of a large proportion of reads being identified as plastid DNA, which were removed from the dataset. Despite this, we included bacterial data in our study because sampling curves indicate that we were able to identify the majority of bacterial taxa present in samples (Supplementary material, Fig. S2). Moreover, previous characterization of microbial communities associated with flowers has frequently observed low species richness (Vannette 2020).

### Landscape context

Land cover within a 1 km buffer of each study orchard was classified into three habitat types: (i) pear orchard, (ii) other fruit orchard (apple and cherry), and (iii) forest. These classifications were performed using the cropland data layer spatial product (USDA 2018). Across our study region, pears were the dominant agricultural crop, although the habitat around individual study sites varied widely from 2 to 66% pear orchards. Other fruit crops had less variability, with 0 to 6% of surrounding land cover, while forest land was highly variable and ranged from 0 to 46%. Forest patches were primarily composed of evergreen trees.

To assess the role of abiotic factors, high resolution climatic metrics for each site were obtained from publicly accessible PRISM data in April 2018. PRISM data is collected at a spatial resolution of 2.5 arcmin (~4km). PRISM data used included elevation, min and max temperature, min and mix vapor pressure deficit (VPD) and precipitation. Vapor pressure deficit is the difference between the amount of moisture in the air and how much moisture the air can hold when saturated, where high VPD indicates drier conditions. As with land cover, the abiotic conditions where sites were located were variable, with elevation ranging from 1152 to 1526 m above sea level, April precipitation ranging from 4.2 to 5.3 cm, minimum temperatures ranging from 2.4 to 3.7 °C, and maximum temperature ranging from 13.6 to 15.7 °C.

### Statistical analyses

We used multivariate linear regression to assess effects of land cover, orchard management, and climate on alpha diversity (Shannon diversity index) and dominance of pear-flower microbiomes. All analyses were conducted using R v. 3.6.1 (R Core Team 2013). To reduce multicollinearity among predictors, we calculated variance inflation factors (VIFs) and used to a threshold of 10 to eliminate variables with problematic covariance. This eliminated temperature, precipitation, and elevation from the alpha diversity models. We calculated multi-model average coefficients based on the 90% confidence interval of top models as well as the importance of each coefficient, which indicated the number of top models in which it appeared.

We also assessed effects of landscape, climate, and farm management on the dominance (relative abundance) of a few focal genera that are highly important for pre- and post-harvest diseases of pear—including putative pathogens and beneficial taxa. These included fungal genera *Aureobasidium, Botrytis*, *Cladosporium*, *Monilinia*, *Mycosphaerella*, *Penicillium*, and beneficial bacteria including *Bacillus, Pantoea*, and *Pseudomonas*. One ASV (BactSeq29) identified as an *Erwinia* sp. was detected at a single orchard in our survey. Given such limited detection, we were unable to perform an analysis of links between variables of interest and *Erwinia* presence and abundance. However, to examine associations between microbial genera and predictors described earlier, we used canonical correlation analysis (CCA), an extension of linear regression that finds linear relationships between combinations of explanatory and response variables which maximize the correlation. Separate models were run on fungi and bacteria of interest.

Differences in species composition among sites could be affected by processes including substitution of taxa, or variation in abundance of particular taxa, so we further evaluated the effects of farm management, land cover, and climate variables on abundance-related and taxa-related aspects of community turnover (microbial beta diversity), and the overall community dissimilarity (which incorporates both processes). Beta diversity was partitioned into abundance-related and taxa-related components of Bray-Curtis dissimilarity using the ‘bray.part’ function in the ‘betapart’ R package. The influence of explanatory variables on these two components of community turnover between sites, as well as their cumulative overall Bray-Curtis dissimilarity was investigated using Restricted Distance-based Analysis (RDA) and AIC model selection, executed using the ‘capscale’ and ‘ordiR2step’ functions in the ‘vegan’ R package. The variance explained by factors included in the top AIC selected models are included in the results.

## ACKNOWLEDGMENTS

We are especially grateful to the growers who allowed us to sample on their properties, and DFJ Sauza for assistance with data analysis. This work was funded by USDA NIFA grants (2014-51106-22096, DW; 2017-67012-26104, RS), Western SARE (SW18-031; DW and RS), the WSDA (#K1986; SD), and the WSU BioAg program.

## REFERENCES

Agrios, G.N. (2005) Plant Pathology, Academic press.

Aizenberg-Gershtein, Y., Izhaki, I., and Halpern, M. (2013) Do honeybees shape the bacterial community composition in floral nectar? PLOS ONE 8: e67556.

Bell T, Tylianakis JM (2016) Microbes in the anthropocene: spillover of agriculturally selected bacteria and their impact on natural ecosystems. Proc Royal Society London Series B 283: 20160896.

Berg, M. and Koskella, B. (2018) Nutrient-and dose-dependent microbiome-mediated protection against a plant pathogen. Curr Biol 28: 2487–2492. e3.

Bulger, M.A., Ellis, M.A., and Madden, L.V. (1987) Influence of temperature and wetness duration on infection of strawberry flowers by *Botrytis cinerea* and disease incidence of fruit originating from infected flowers. Phytopathology 77: 1225–1230.

Busby, P.E., Soman, C., Wagner, M.R., Friesen, M.L., Kremer, J., Bennett, A., et al. (2017) Research priorities for harnessing plant microbiomes in sustainable agriculture. PLOS Biol 15:e2001793.

Callahan, B.J., McMurdie, P.J., Rosen, M.J., Han, A.W., Johnson, A.J.A., and Holmes, S.P. (2016) DADA2: high-resolution sample inference from Illumina amplicon data. Nat Methods 13: 581–583.

Davis, N.M., Proctor, D.M., Holmes, S.P., Relman, D.A., and Callahan, B.J. (2018) Simple statistical identification and removal of contaminant sequences in marker-gene and metagenomics data. Microbiome 6: 226.

DuPont, S.T. and John Strohm, C. (2020) Integrated pest management programmes increase natural enemies of pear psylla in Central Washington pear orchards. J Appl Entomol 144: 109–122.

Edwards, J., Johnson, C., Santos-Medellín, C., Lurie, E., Podishetty, N.K., Bhatnagar, S., et al. (2015) Structure, variation, and assembly of the root-associated microbiomes of rice. Proc Natl Acad Sci 112: E911–E920.

Fahrig, L., Baudry, J., Brotons, L., Burel, F.G., Crist, T.O., Fuller, R.J., et al. (2011) Functional landscape heterogeneity and animal biodiversity in agricultural landscapes. Ecol Lett 14: 101–112.

Fahrig, L. and Nuttle, W.K. (2005) Population ecology in spatially heterogeneous environments. In Ecosystem function in heterogeneous landscapes. Springer, pp. 95–118.

Grady, K.L., Sorensen, J.W., Stopnisek, N., Guittar, J., and Shade, A. (2019) Assembly and seasonality of core phyllosphere microbiota on perennial biofuel crops. Nat Commun 10: 1–10.

Johnson, K.B. and Stockwell, V.O. (1998) Management of fire blight: a case study in microbial ecology. Annu Rev Phytopathol 36: 227–248.

Johnson, K.B. and Temple, T.N. (2013) Evaluation of strategies for fire blight control in organic pome fruit without antibiotics. Plant Dis 97: 402–409.

Klein, A.-M., Brittain, C., Hendrix, S.D., Thorp, R., Williams, N., and Kremen, C. (2012) Wild pollination services to California almond rely on semi-natural habitat. J Appl Ecol no-no.

de Kock, S.L. and Holz, G. (1992) Blossom–end rot of pears: Systemic infection of flowers and immature fruit by *Botrytis cinerea*. J Phytopathol 135: 317–327.

Lindow, S.E. and Andersen, G.L. (1996) Influence of immigration on epiphytic bacterial populations on navel orange leaves. Appl Environ Microbiol 62: 2978–2987.

Logan, J., Mueller, M.A., and Searcy, M.J. (2000) Microclimates, peach bud phenology, and freeze risks in a topographically diverse orchard. HortTechnology 10: 337–340.

Lopez, G. and DeJong, T.M. (2007) Spring temperatures have a major effect on early stages of peach fruit growth. J Hortic Sci Biotechnol 82: 507–512.

Lymperopoulou, D.S., Adams, R.I., and Lindow, S.E. (2016) Contribution of vegetation to the microbial composition of nearby outdoor air. Appl Environ Microbiol 82: 3822–3833.

McGhee, G.C. and Sundin, G.W. (2011) Evaluation of kasugamycin for fire blight management, effect on nontarget bacteria, and assessment of kasugamycin resistance potential in Erwinia amylovora. Phytopathology 101: 192–204.

McNicol, R.J., Williamson, B., and Dolan, A. (1985) Infection of red raspberry styles and carpels by Botrytis cinerea and its possible role in post-harvest grey mould. Ann Appl Biol 106: 49–53.

Mueller, U.G. and Sachs, J.L. (2015) Engineering microbiomes to improve plant and animal health. Trends Microbiol 23: 606–617.

Muñoz, M., Faust, J.E., and Schnabel, G. (2019) Characterization of *Botrytis cinerea* from commercial cut flower roses. Plant Dis 103: 1577–1583.

Ngugi, H.K. and Scherm, H. (2006) Biology of flower-infecting fungi. Annu Rev Phytopathol 44:261–282.

Ngugi, H.K. and Scherm, H. (2004) Pollen mimicry during infection of blueberry flowers by conidia of *Monilinia vaccinii-corymbosi*. Physiol Mol Plant Pathol 64: 113–123.

Petrasch, S., Knapp, S.J., Van Kan, J.A., and Blanco-Ulate, B. (2019) Grey mould of strawberry, a devastating disease caused by the ubiquitous necrotrophic fungal pathogen *Botrytis cinerea*. Mol Plant Pathol 20: 877–892.

Pii, Y., Mimmo, T., Tomasi, N., Terzano, R., Cesco, S., and Crecchio, C. (2015) Microbial interactions in the rhizosphere: beneficial influences of plant growth-promoting rhizobacteria on nutrient acquisition process. A review. Biol Fertil Soils 51: 403–415.

Pusey, P.L., Stockwell, V.O., and Mazzola, M. (2009) Epiphytic bacteria and yeasts on apple blossoms and their potential as antagonists of *Erwinia amylovora*. Phytopathology 99: 571–581.

R Core Team (2013) R: A Language and Environment for Statistical Computing, Vienna, Austria: R Foundation for Statistical Computing.

Schaeffer, R.N., Vannette, R.L., Brittain, C., Williams, N.M., and Fukami, T. (2017) Non-target effects of fungicides on nectar-inhabiting fungi of almond flowers. Environ Microbiol Rep 9: 79–84.

Shemshura, O., Alimzhanova, M., Ismailova, E., Molzhigitova, A., Daugaliyeva, S., and Sadanov, A. (2020) Antagonistic activity and mechanism of a novel Bacillus amyloliquefaciens MB40 strain against fire blight. J Plant Pathol 1–9.

Smessaert, J., Van Geel, M., Verreth, C., Crauwels, S., Honnay, O., Keulemans, W., and Lievens, B. (2019) Temporal and spatial variation in bacterial communities of “Jonagold” apple (*Malus* x *domestica* Borkh.) and “Conference” pear (*Pyrus communis* L.) floral nectar. MicrobiologyOpen 8: e918.

Smith, T.J. (2000) Overview of tree fruit production in the Pacific Northwest United States of America and southern British Columbia, Canada. In IV International Symposium on Mineral Nutrition of Deciduous Fruit Crops 564. pp. 25–30.

Smith, T.J., and P. L. Pusey. 2010. CougarBlight 2010 ver. 5.0: a significant update of the CougarBlight fire blight infection risk mode. Acta Horticulturae 896:331–338.

Smith OM, Cohen AL, Reganold JP, Jones MS, Orpet RJ, Taylor JM, Thurman JH, Cornell KA, Olsson RL, Ge Y, Kennedy CM, Crowder DW (2020) Landscape context affects the sustainability of organic farming systems. PNAS 117, 2870–2878.

Stockwell, V.O., Johnson, K.B., Sugar, D., and Loper, J.E. (2002) Antibiosis contributes to biological control of fire blight by *Pantoea agglomerans* strain Eh252 in orchards. Phytopathology 92: 1202–1209.

Stockwell, V.O., McLaughlin, R.J., Henkels, M.D., Loper, J.E., Sugar, D., and Roberts, R.G. (1999) Epiphytic colonization of pear stigmas and hypanthia by bacteria during primary bloom. Phytopathology 89: 1162–1168.

Sundin, G.W., Werner, N.A., Yoder, K.S., and Aldwinckle, H.S. (2009) Field evaluation of biological control of fire blight in the eastern United States. Plant Dis 93: 386–394.

Toju, H., Peay, K.G., Yamamichi, M., Narisawa, K., Hiruma, K., Naito, K., et al. (2018) Core microbiomes for sustainable agroecosystems. Nat Plants 4: 247.

Vannette, R.L. (2020) The floral microbiome: Plant, pollinator, and microbial perspectives. Annu Rev Ecol Evol Syst 51:.

Vannette, R.L. and Fukami, T. (2017) Dispersal enhances beta diversity in nectar microbes. Ecol Lett 20: 901–910.

Vannier, N., Agler, M., and Hacquard, S. (2019) Microbiota-mediated disease resistance in plants. PLoS Pathog 15: e1007740.

Vurukonda, S.S.K.P., Vardharajula, S., Shrivastava, M., and SkZ, A. (2016) Enhancement of drought stress tolerance in crops by plant growth promoting rhizobacteria. Microbiol Res 184: 13–24.

Wang, Q., Garrity, G.M., Tiedje, J.M., and Cole, J.R. (2007) Naive Bayesian classifier for rapid assignment of rRNA sequences into the new bacterial taxonomy. Appl Environ Microbiol 73:5261–5267.

