## Supplementary figures and images for "Orchard Management and Landscape Context Mediate the Floral Microbiome of Pear"

### Supplemental Figure 1

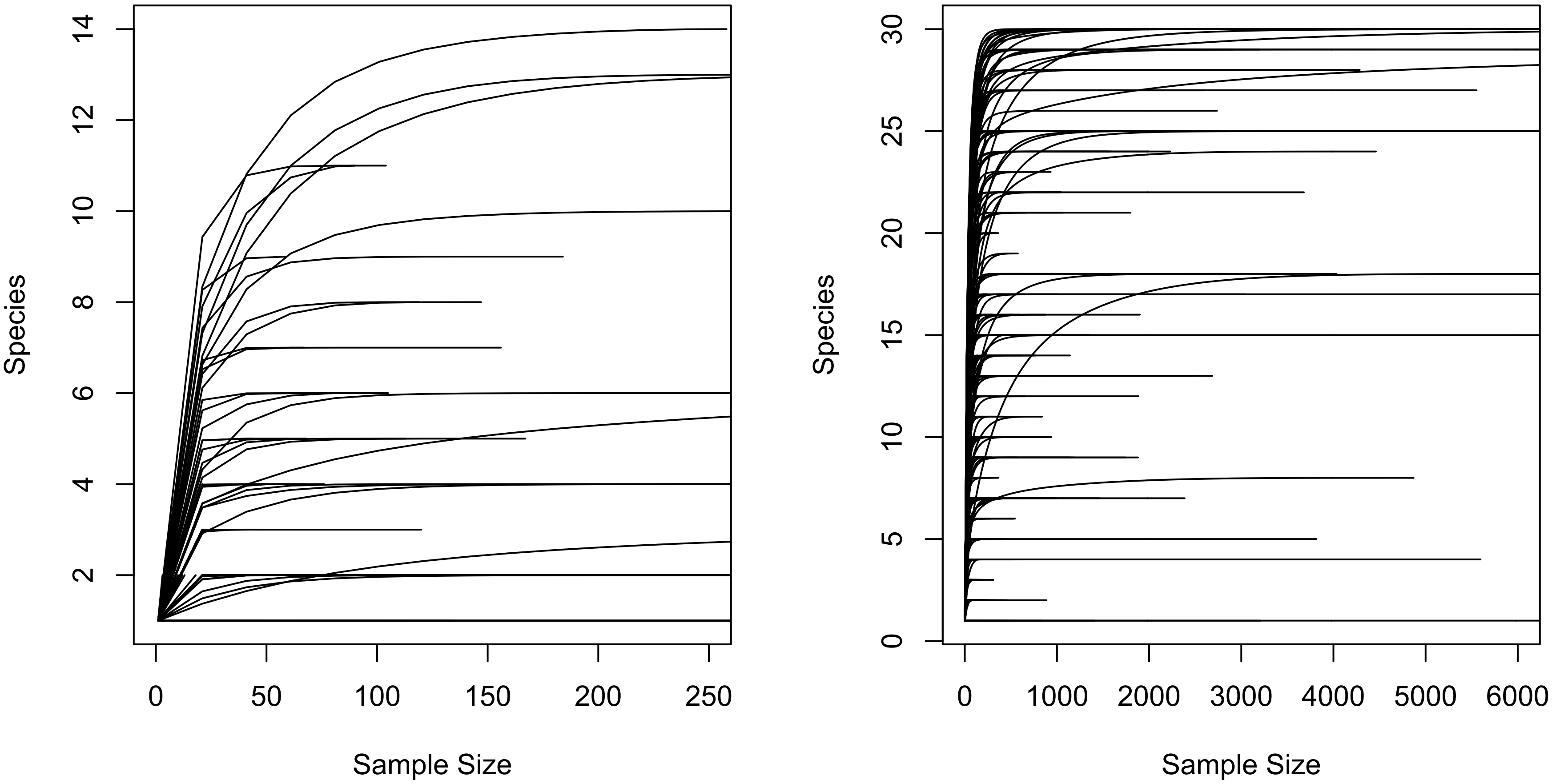

### Supplemental Figure 2

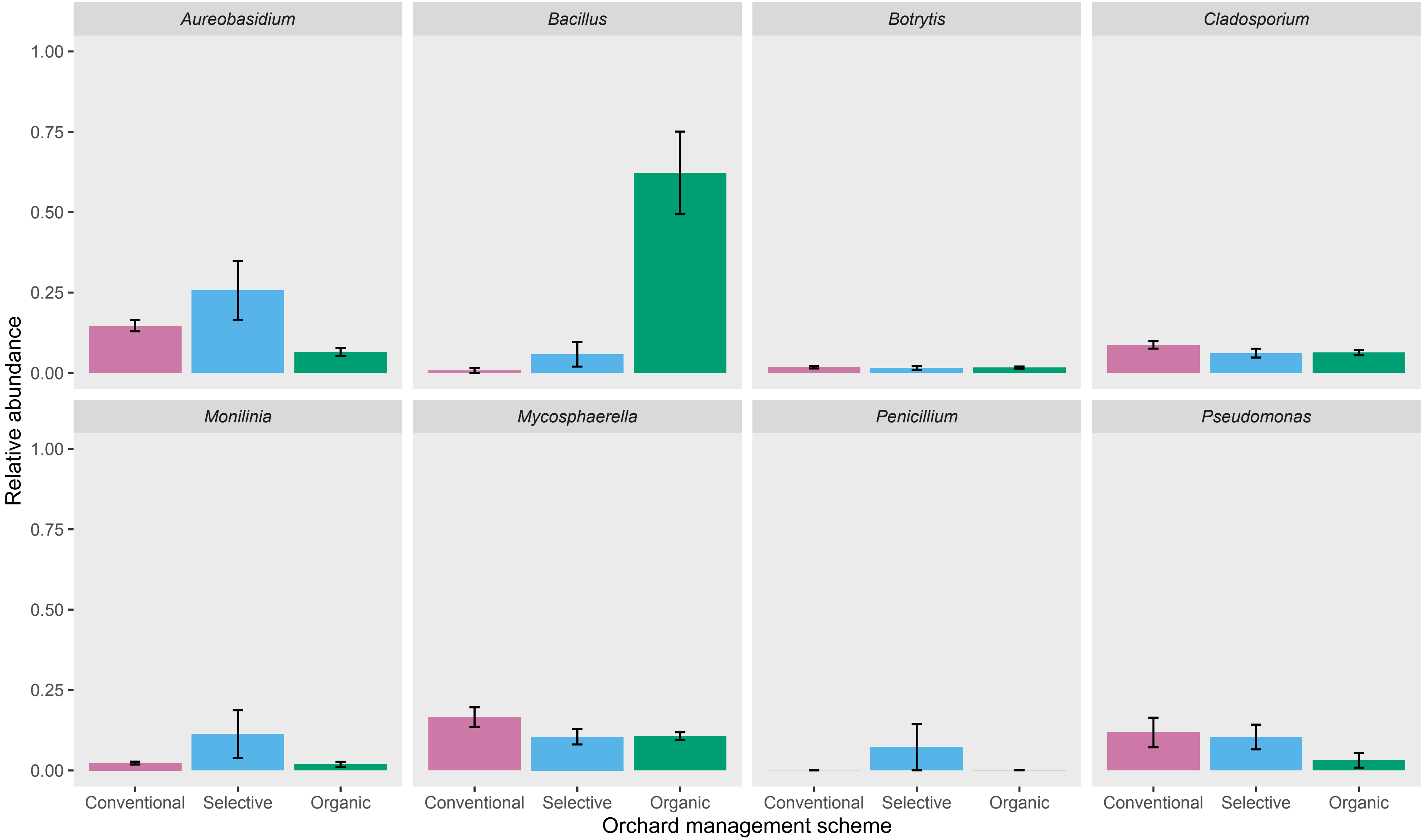
